# The genomic architecture of flowering time varies across space and time in *Mimulus guttatus*

**DOI:** 10.1101/111203

**Authors:** Patrick J. Monnahan, John K. Kelly

## Abstract

The degree to which genomic architecture varies across space and time is central to the evolution of genomes in response to natural selection. Bulked-segregant mapping combined with pooled sequencing provides an efficient method to estimate the effect of genetic variants on quantitative traits. We develop a novel likelihood framework to identify segregating variation within multiple populations and generations while accommodating estimation error on a sample- and SNP-specific basis. We use this method to map loci for flowering time within natural populations of *Mimulus guttatus*, collecting the early and late flowering plants from each of three neighboring populations and two consecutive generations. We find appreciable variation in genetic effects on flowering time across both time and space; the greatest differences evident between populations. Structural variants, such as inversions, and genes from multiple flowering time pathways exhibit the strongest associations with flowering time. It is also clear that genotype-by-environment interactions are an important influence on flowering time variation.

## Introduction

The standing genetic variation in a population is the raw material for evolution. For quantitative traits, a basic question is whether the architecture of this variation is consistent across populations of a species, or even within a single population through time. Consistency requires not only for the same polymorphisms to be present in each population, but also that the genotype-to-phenotype mapping is stable across space and time. The consistency of genomic architecture is relevant to many outstanding questions: How general are the results from QTL mapping studies, typically done on a single population evaluated in a single environment? How frequently will parallel selection pressures produce parallel genetic changes (Cohan 1984a; Cohan 1984b; Colosimo *et al.* 2005; Monnahan and kelly 2015b)? How influential are factors such genotype-by-environment (GxE) interactions in generating inconsistent architecture from spatial and temporal environmental variation? To what extent do such factors alter the balance of evolutionary forces that maintain the quantitative trait variation in the first place?

To address the question of consistency, we performed a bulked-segregant mapping experiment of flowering time variation across multiple population of *Mimulus guttatus* over two generations in the field. Bulked-segregant mapping (Michelmore *et al.* 1991) identifies loci that are divergent between the tails of the distribution of a phenotype, in this case the earliest and latest flowering plants in a population. Quantitative Trait Loci (QTL) for flowering time should exhibit allele frequency divergence between groups (bulks). Because the selection of bulks is equivalent to a single generation of (bi-directional) truncation selection, the expected magnitude of this difference is directly proportional to the “average effect” of alleles on the trait (Fisher 1941; Latter 1965; Kimura and Crow 1978). The average effect measures the association between alleles and phenotypes (Falconer and Mackay 1996), and the extent to which the average effect changes with context directly assays importance of that context on variation. Changes in average effect across environments estimates the effect of genotype-by-environment interaction. Changes in average effect owing to different genetic backgrounds estimate the effect of epistasis.

The three populations chosen for this study are geographically proximal (within 7km of each other in the Central Cascades of Oregon, U.S.A) and exhibit extensive shared polymorphism (Monnahan *et al.* 2015). When polymorphisms are shared, differences among populations must be due to difference in the expression of variation rather than the presence or absence of different alleles. The change in allele frequency between early and late flowering individuals within a population (which we call Δp_EL_) can differ between populations for numerous reasons. If the mapping from genotype to phenotype is constant, Δp_EL_ will differ if the allele frequency is intermediate in one population but extreme in the other. Without differences in allele frequency, Δp_EL_ will differ if the distribution of genetic backgrounds differs between the populations and that influences expression of the focal locus. Environmental differences among populations can alter the magnitude or even direction of Δp_EL_. Furthermore, GxE can generate heterogeneity in ΔpEL between generations within a population if there are temporal changes in the environment.

Our trait of study, flowering time, is typically highly polygenic and responsive to numerous environmental variables (Bernier and Périlleux 2005; Wellmer and Riechmann 2010; Blümel *et al.* 2015). It is central to numerous ecological and evolutionary processes. For many plants, the time of flower production is a major determinant of fitness because access to pollination and resources necessary to complete reproduction (set seed) vary over the course of a growing season. This is particularly true for annual *M. guttatus*, in which plants must flower and set seed before the cessation of water availability. Although late-flowering plants tend to produce more seed than their early-flowering counterparts, they risk desiccation prior to seed set (Mojica and Kelly 2010; Mojica *et al*. 2012). This tradeoff may be relevant to the maintenance of genetic variation in flowering time and will surely play a role in how these populations evolve in response to a changing climate. Shifts in flowering time due to climate change have already been observed for a number of species (Fitter and Fitter 2002).

### Measuring genetic effects

Estimating the contribution of individual loci to quantitative trait variation is a challenge (McCarthy *et al*. 2008; King *et al*. 2012). Genetic effects may be subtle and thus difficult to distinguish from random fluctuations. In Bulked-Segregant mapping, differences in allele frequency owing to random sampling should usually be small if bulks are large, but occasional, large random fluctuations are inevitable. In the present study, statistical difficulties are acute given that we wish not only to detect loci affecting a trait, but also test whether these effects vary with year or population. To this end, we develop a likelihood-based hypothesis testing framework analogous to the factorial Analysis of Variance, in which we can test for marginal effects as well as interactions between factors. The marginal (average) effect of a locus on flowering time is important, but we also wish to know whether differences between bulks change with population or year. A locus with variable effects across levels of these other factors should inflate the “interaction” test statistic.

We used pooled population sequencing, “PoolSeq” (Schlötterer *et al*. 2014), to estimate allele frequencies in each bulk throughout the genome. Each bulk makes a single pool of DNA to be sequenced with the resulting read counts estimating allele frequencies. This method has been applied successfully to study population differentiation (Fabian *et al.* 2012), population dynamics of transposable elements (Kofler *et al.* 2012), and the genomic response to selection (Turner *et al.* 2011; Kelly *et al.* 2013; Beissinger *et al.* 2014; Tobler *et al.* 2014). An important difficulty is accommodating the variance introduced by the sampling events prior to sequencing. These include, but are not limited to, sampling of individuals from populations, sampling DNA into pools, sampling events during library preparation (particularly, PCR), and sampling of fragments for sequencing. Multiple methods have been proposed to estimate the variance in allele frequency estimates obtained from PoolSeq data (Magwene *et al.* 2011; Gautier *et al.* 2013; Kelly *et al.* 2013; Lynch *et al.* 2014). Here, we build on a method based on Fisher’s angular transformation of allele frequency (Fisher and Ford 1947) using a robust estimator for the variance of dispersive processes (Kelly *et al.* 2013).

In addition to the genome-wide mapping, we estimate flowering time effects for five structural variants (chromosomal inversions) found to be segregating in one or more of the populations. These variants, identified in prior mapping studies (Fishman and Saunders 2008; Lowry and Willis 2010; Holeski *et al.* 2014; Lee *et al.* 2016), are located on chromosomes 5, 6, 8, 10, and 11. Previous studies have demonstrated phenotypic effects, including developmental timing, for three of these loci (*inv6, inv8*, and *D*). The present study provides further evidence of natural selection on alternative orientations of the inversions. Also, the inclusion of “known loci” provides important ground-truthing for genome scans in which the overwhelming majority of SNPs are effectively anonymous.

Considering both SNPs and structural variants, this study provides several striking observations regarding genomic variation for flowering time in natural populations of *M. guttatus.* Depending on the population and year, we find anywhere from tens to thousands of SNPs that differ in frequency between early and late flowering plants, broadly distributed throughout the genome. Although individual SNPs are almost entirely idiosyncratic with regard to significance, there is appreciable overlap in the genomic regions harboring this variation. Furthermore, we find that the extent of variability over time itself varies between populations. The Quarry population, a recently established annual/perennial hybrid swarm, exhibits many more early-late divergent SNPs compared to the other two, and the allele frequency divergence at these SNPs tends to be much more consistent across years. In the following sections, we describe our likelihood framework in detail and interpret the results in relation to the expected degree and scale of parallel evolution, as well as the generality of genetic mapping studies.

## Theory

In this section, we describe a likelihood framework for testing divergence in allele frequency; first between two bulks (Early vs. Late) and then extended to treat multiple contrasts simultaneously. Following Fisher and Ford (1947), we conduct tests on transformed allele frequencies: *x̂* = 2 arcsin 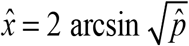 where *ρ̂* is the estimated allele frequency (fraction of reads bearing the specified base at a SNP) in a bulk. Transformed allele frequency can be treated as normally distributed values with a variance determined by the series of sampling events that ultimately produce the observed read counts. These events contribute additively and in a reciprocal manner to the sampling variance (Kelly *et al.* 2013). For a single bulk, 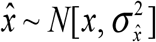 where x is the true (transformed) allele frequency and 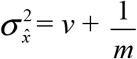. Here, *v* is a bulk-specific variance that aggregates the effects of sampling of individuals into bulks, sampling of DNA into the pooled sample, and PCR sampling during library preparation, and *m* is read depth at a SNP. *v* is common to all SNPs in a the bulk while the read depth will vary among SNPs. The simple null hypothesis that Δp_EL_ = 0 (allele frequency is the same across bulks) is evaluated with transformed allele frequencies as:

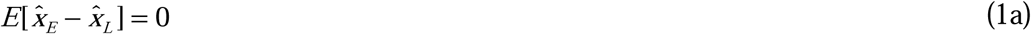

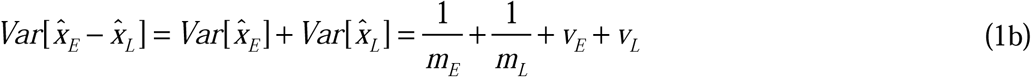

Given values for *m_E_*, *m_L_*, *v_E_*, and *v_L_*, we can calculate the likelihood of any observed difference from the normal density function. The read depths in a sample are directly observed while the *v* terms are estimated from a genome-wide aggregation of data (procedure described below).

A likelihood ratio test statistic (LRT) for a difference between bulks requires a maximum likelihood estimator for the common allele frequency (same in each bulk):

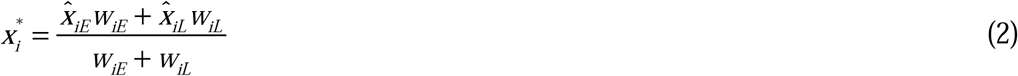

Here, iE *x̂* and *x̂*_iL_ are the estimates from each bulk at site *i* and *w* terms are the reciprocal of the bulk/site-specific variances. The log-likelihood of the data under the null model is:

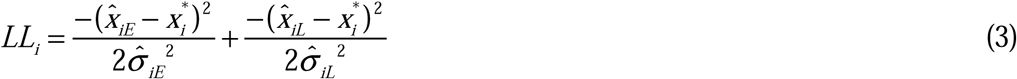

Here, 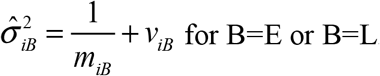 for B=E or B=L. This can be compared to an unconstrained model, where *x̂_iE_* ≠ *X_iL_*, with a separate mean estimated for each bulk. Since there is only one observation (allele frequency) in each sample, the estimate is simply the observation, and the log-likelihood of the unconstrained model becomes zero. The LRT is then −2 times equation (3), and a p-value for the test is obtained from a chi-square distribution with 1 degree of freedom.

These calculations can be generalized to consider two contrasts (Δp_EL_ from different populations or generations) simultaneously. Table 1 outlines three models appropriate to test for heterogeneity of such contrasts. These models are nested: M1 is a special case of M2, M2 a special case of M3. Comparing two generations within a population, a significant LRT for M1 vs. M2 indicates a *marginal* effect (average divergence) between bulks across the two generations. A significant test for M2 vs. M3 indicates heterogeneous divergence, Δp_EL_ differs between generations (i.e. there is an *interaction* between generation and bulk divergence). Contrasting M1 v. M3 represents an overall test for significant divergence across both years because it is the sum of the two LRTs from the former tests. The degrees of freedom are 1 for the former tests, and 2 for the latter.

**Table 1.** Models to establish significance of marginal and heterogeneous genetic effects across the two contrasts (between years or between populations) within a context.

| Model | Description | Parameter Constraints |
| --- | --- | --- |
| M1 | No difference between E/L | $x_{E1} = x_{L1}; x_{E2} = x_{L2}$ |
| M2 | $\Delta p_{EL}$ consistent | $x_{E1} = x_{L1} + \alpha;$<br>$x_{E2} = x_{L2} + \alpha$ |
| M3 | $\Delta p_{EL}$ variable | $x_{E1} \neq x_{L1}; x_{E2} \neq x_{L2}$ |

Calculating the likelihoods of M1 and M3 requires no additional derivations beyond equations (2)-(3) as each is a sum of the log-likelihoods for each population. For M2, the MLEs for the 3 parameters are:

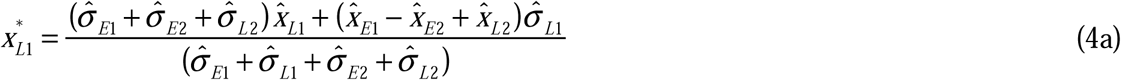

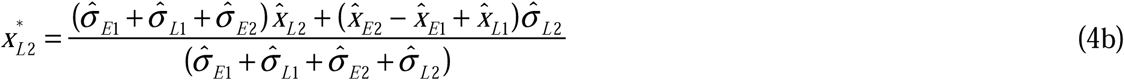

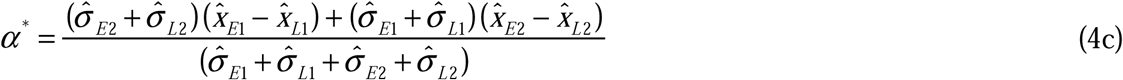
where 1 and 2 simply designate the population (or generation) being considered. The log-likelihood of the data for M2 is:

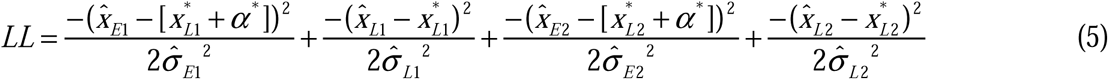

Figure 1 illustrates some key features regarding the marginal and interaction tests as applied to data such as ours (typical read depths and bulk-specific variances). As expected, if the observed Δ*p* is the same in both populations (or generations), the LRT for an interaction test (M2 v M3) is 0. When the opposite is true (equal magnitude Δ*p*, but different sign), the LRT for the marginal effect test (M1 v M2) is zero (note that, when the results for negative Δ*p*_2_ are plotted instead of the positive values displayed in Figure 1, the solid line follows the dotted line trajectory and vice versa). When Δ*p* is non-zero in only one population (or generation), both tests are equally powered (the dashed and solid *black* line are perfectly overlapping when Δ*p*_2_ = 0). As observed *average* Δ*p* increases, so does the LRT for marginal effects, regardless of whether Δ*p*_1_ ≠ Δ*p*_2_. The interaction LRT increases as Δ*p*_1_ and Δ*p*_2_ diverge. For a fixed difference between Δ*p*_1_ and Δ*p*_2_, the interaction LRT increases as *average* Δ*p* increases. For example, the interaction LRT is 7.08 when Δ*p*_1_ is 0.75 and Δ*p*_2_ is 0.25 (i.e. divergence between Δ*p*_1_ and Δ*p*_2_ is 0.5), but is much higher (21.93) when Δ*p*_1_ is 1.0 and Δ*p*_2_ 0.5. However, heterogeneity is necessarily limited for very large Δ*p* (since each population has a maximum value of 1) unless the direction of effect differs between populations. For polygenic traits, Δ*p* nearing 1 or −1 is unlikely except when greatly exaggerated by sampling. For example, a locus that exhibits an additive effect of 0.5 phenotypic standard deviations (2a = 1.0), has an expected Δ*p* of 0.44 (if *p* = 0.5), although the observed Δ*p* can be greater owing to chance.

**Figure 1.**
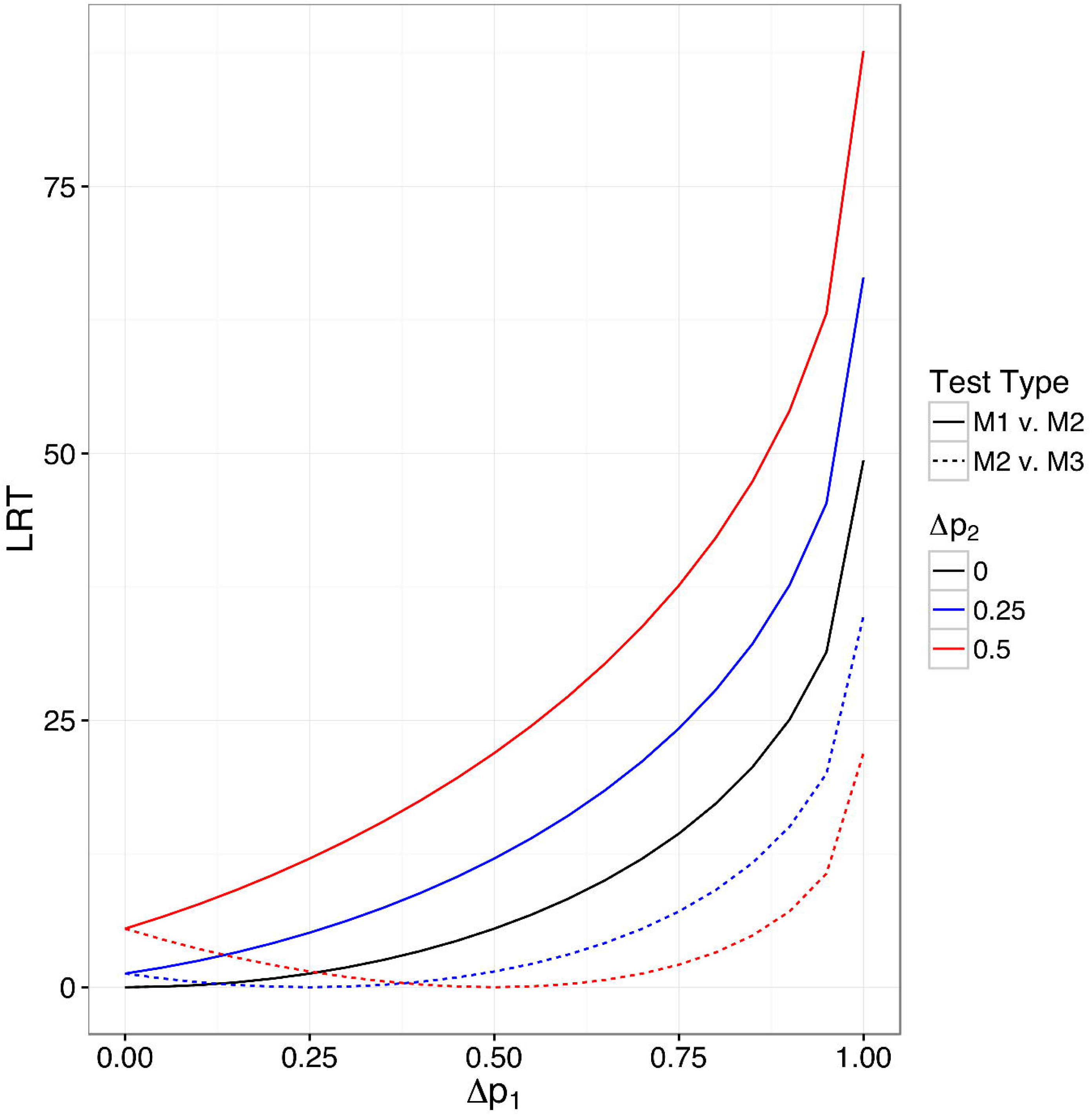
Likelihood Ratio Test values for the consistent effect (M1 v M2) and interaction (M2 v M3) test as a function of allele frequency difference in the two populations. Note that solid and dashed lines are perfectly overlapping for Δ*p*_2_ *=* 0. In all cases, 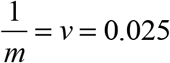 for both populations in the contrast.

### Simulations

We developed a simulation framework to confirm the behavior of our testing procedures under different scenarios, using the real data to calibrate these simulations. We average the observed allele frequencies in the two samples (e.g. Early and Late) to set *p* for each population/year and incorporate sampling error using observed read counts and *v* terms. We simulated new values for each site and sample by adding to the population/year allele frequency a deviation due to sampling error as well as a deviation due to an effect (*a*) of that site. For the sampling error deviation, we add a value drawn from 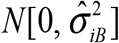, recalling that the sampling variance at a site, 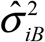, is the sample-specific variance (*v*) plus 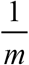. We first investigated the behavior of our testing procedure under a purely neutral scenario (i.e. *a* = 0 for all sites). These simulations confirm that our Likelihood Ratio Tests follow the predicted null distributions (chi-square with 1 d.f. for M1 vs M2, chi-square with 2 d.f. for M1 vs M3) if a SNP is neutral (no effect on phenotype).

Next, we consider scenarios where a subset of SNPs exhibit a constant effect on Δp_EL_, to provide a baseline for comparison of observed heterogeneity in ΔpEL. Given that sampling bulks is a form of truncation selection, the expected allele frequency difference between early and late flowering plants can be calculated given values for the effect size (*a*) and the intensity of selection (*i*):

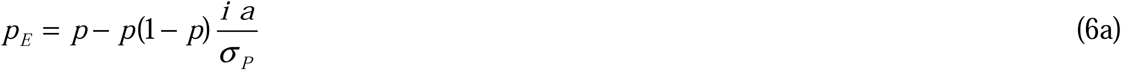

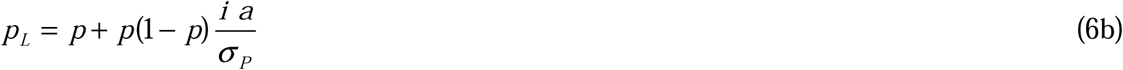

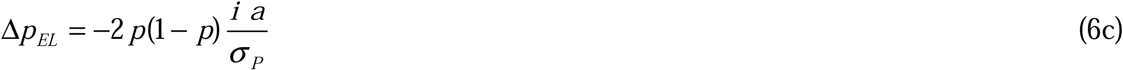

Where *p* is the overall frequency in the population (Falconer and Mackay (1996), ch 11). The intensity of selection was determined using a truncated normal distribution in which 10% of individuals exceed the truncation point (*i* = 1.755). This was based on approximations of population size during sampling periods relative to full bloom. We grossly approximate the distribution of standardized allelic effects, 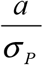, by assuming that some fraction of sites are neutral with respect to flowering time (1 - *f_0_*) and have *a* = 0, while the remaining sites have nonzero effect of constant magnitude, *c* (the sign of *c* is chosen randomly for each site). For each of the four contexts in which we investigated heterogeneity (IM, Q, 2013, and 2014), we performed a heuristic search for values of *f_0_* and *c* that generate a distribution of LRT for the marginal effect test (M1 v. M2) that closely matches the distributions from the real data. Our matching criteria is based on the observed proportion of sites exceeding specified values of LRT, in this case Pr[LRT>10] and Pr[LRT>15]. Here, we aim to match the tails of the LRT distribution as this information pertains most directly to *f_0_* and *c* (given that *f_0_* is likely small).

To accumulate this information into a single measure (*Z_diff_*) we sum the standardized difference between simulated and real data.

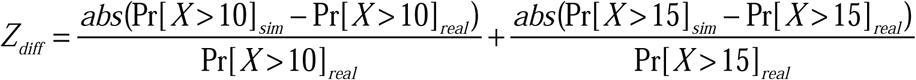

## Materials and Methods

### Plant Collection and phenotyping

The three populations are located in the central Oregon cascades: Quarry (44.3454243 N, 122.1362023 W), IM (44.402217 N, 122.153317 W), and BR (44.373238 N, 122.130675 W) and described in detail in (Monnahan *et al*. 2015). In a particular population and year, we sampled early and late flowering plants according to the following scheme. First, we established several parallel transects, perpendicular to the slope of the hillside, totaling ~30 m. We then divided each transect into 30 cm intervals. We chose sampling times based on density of *flowering* plants along a transect. For the early flowering plants, we sampled a transect as soon as 2 *flowering* plants could be found within ~15 cm on either side of the transect within each 30 cm interval. We estimate this to correspond to about 5–10% of the total population. For the late flowering plants, we waited until plants density was similar to the early sampling event. If a particular interval along a transect had several flowering plants within 15 cm on each side, we randomly selected the two plants nearest the transect line. Whole plants were collected and stored in dry ice until frozen at −20° C. Since collection times were dependent on density of flowering plants, sample times varied across populations and across years (see Supplemental Table 1 for collection dates). Early bulks were collected earlier in 2014 for all populations. The late bulk for Q was collected earlier in 2014. There was a very hot and dry spell that wiped out the BR population shortly after collecting the Early bulk; therefore, we did not perform an Early/Late contrast for Browder Ridge in 2014.

### Sequencing and SNP calling

We extracted DNA from each individual collected over 2013 and 2014. Individuals’ DNA was pooled in equal amounts corresponding to the year, population, and bulk in which they were collected. Each pooled sample was whole-genome sequenced on an Illumina HiSeq 2500 (paired-end 100bp reads). The 2013 samples were sequenced in three High-Output lanes, while the 2014 samples were sequenced with two High-Output lanes. Two additional lanes (Rapid-Runs) were performed in order to equilibrate coverage across samples. We combined data from all lanes to create 10 fastq sets corresponding to each of the sampling bulks and ran Scythe (https://github.com/vsbuffalo/scythe) and Sickle (Joshi and Fass 2011) for each fastq file to remove adaptors and trim low quality sites, respectively. We mapped reads to the *M. guttatus* v2 genome build using BWA and removed PCR duplicates using Picard Tools. We then called SNPs using the GATK UnifiedGenotyper with the down-sampling feature suppressed (‘-dt NONE’). The read counts in the variant call file corresponding to each of the sampling bulks are the input for subsequent likelihood analyses. A SNP was included for Δp_EL_ within a sample only if read depth per bulk was 25-100 reads and allele frequency (both bulks combined) was between 0.05 and 0.95. We imposed the upper bound of 100 reads to exclude paralogous mappings.

### Estimation of v terms

In equations (1)-(5), the bulk-specific variance terms (*v*) are treated as known constants. Prior to hypothesis testing, we estimate these variances using a procedure similar to that in Kelly *et al*. (2013). We first perform a series of pairwise contrasts (difference in transformed allele frequencies at each site) between the four bulks within a population (6 pairwise contrasts for IM and Q; 3 for BR). Under the assumption that divergence among the bulks will be random for most of the genome, each of these contrasts will be centered on zero with a variance equal to the sum of the individual sample variances (i.e. the two *v* terms plus each sample’s variance due to read depth). We estimate *Var*[*x̂*_1_ - *x̂*_2_] using the interquartile range (Supplemental Appendix 1) of the genome-wide distribution of *x̂*_1_ - *x̂*_2_, which is robust to the presence of outliers (SNPs that are correlated with flowering time or divergent across generations). We also estimate the read depth variance as the average of 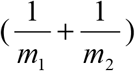 across all SNPs for the contrasted samples (1 and 2). Following equation (1b), the two *v* values are equal to the estimated total variance for the contrast minus the read depth variance. Repeating this entire process for the remaining 5 pair-wise contrasts ultimately produces 6 equations that are a function of 4 unknowns. This system is overdetermined (i.e. there are more equations than unknowns), and so we utilize the method of General Least Squares to obtain an optimal compromise for the *v* terms (Lynch and Walsh 1998) as well as an estimate of their sampling variance. The only additional information necessary to calculate the *v* terms are estimates of the (co)variance of the 6 contrast variances, which we obtain by jackknifing the original data set of read counts, recalculating the contrast variances after deleting a portion (0.1) of the original data. The small sampling variance, and thus standard errors, for the *v* terms justifies treating these values as constants in our analyses and simulations (Supplemental Appendix 1).

### Structural variants

Initial genotyping confirmed that five structural variants (*inv5, inv6, inv8, inv10*, and D) were segregating in one or more of the populations. Two of the variants, *inv6* and the meiotic drive locus (*D*), were previously only known to segregate within IM. The others were mapped in crosses between annual and perennial genotypes of *M. guttatus.* We cannot identify a single diagnostic SNP for any of these features (recognizing alternative orientations from alternative SNP bases). For *inv6* and D, the derived haplotype is associated with a single predominant nucleotide sequence over >4mb, but the ancestral orientation is internally variable. For the other inversions, both alternative orientations harbor many distinct sequences. For each feature however, there are differences in SNP allele frequency between the populations of sequences within each orientation. We thus developed a SNP set that is predictive of orientation for each inversion.

We used a collection of ten fully sequenced inbred lines from the IM population to generate the SNP sets for *inv6* and *D* (Flagel *et al.* 2014; Lee *et al.* 2016). PCR based genotyping of length polymorphic markers indicate that four of the ten lines carry the derived orientation at *D*, while two of ten have the derived orientation for *inv6.* We found 11,848 SNPs for the *Drive* locus in which at least 5/6 of the non-drive lines harbor the alternative base (the *Drive* haplotype is always Reference base because the Reference genome is based on a line homozygous for *Driver*). These SNPs are located within three distinct intervals on chromosome 11 of the genome build (5.7-11.6 Mb, 13.9-14.1 Mb, 16.6-21.1 Mb) because the region is misassembled in the reference genome sequence (Holeski *et al.* 2014). We identified 26,739 SNPs for *inv6* in which the two lines homozygous for the derived orientation are fixed for the alternative base and the other eight lines are fixed for reference in the genomic interval 1.34-7.61 Mb of chromosome 6 (Lee *et al.* 2016).

To develop a predictive SNP set for *inv5, inv8*, and *inv10*, we assembled and interrogated ten whole genome sequences, one plant from each of five annual populations (MAR3, REM8-10, CAC6G, LMC 24, and SLP19) and five perennial populations (TSG3, BOG10, YJS6, SWB, and DUN). All data is available from the Sequence Read Archive (http://www.ncbi.nlm.nih.gov/sra). The relevant genomic regions are 10-18 Mb of chromosome 5 (*inv5*), 1.5-7.0 Mb of chromosome 8 (*inv8*), and 2.0-6.0 Mb of chromosome 10 (*inv10*). To include a SNP in the diagnostic set for a feature, we required that the reference base predominate in annuals and vice versa: At least 3 lines of each type were called and at most one contradiction (annual line is alternative or perennial line is reference) was tolerated. With these conventions, the reference base within a structural variant identifies the derived orientation for *D*, the ancestral orientation for *inv6*, and the annual orientation for the other three loci. We averaged Δp_EL_ across SNPs within a feature to estimate the change in orientation frequencies between Early and Late flowering plants. Because the correlation between SNP alleles (reference vs alternative) and orientation is imperfect, the average SNP Δp_EL_ should ***underestimate*** the magnitude of Δp_EL_ for inversion orientations.

## Results

### Polymorphism

After filtering, we identified approximately 7.5 million SNPs, most of which were segregating in more than one population. However, the pattern of shared polymorphism is asymmetric (Supplemental Figure 1). For SNPs in IM or BR with a minor allele frequency of at least 10%, 94% are segregating in the samples from the other two populations. This is nearly complete overlap given that a population sample with as few as 25 reads is counted (and an allele at ≤10% population frequency will often fail to be sampled). In contrast, Quarry has a higher frequency of intermediate frequency SNPs that are rare or fixed in IM and BR. SNPs in the 10-90% range in Quarry are not evident in other populations about 25% of the time.

### Tests for association with flowering time

We first tested for significant Δ*p_EL_* within each population/year and then performed the structured hypothesis testing of Table 1. For the former, the Quarry population exhibited an order of magnitude greater number of significant sites (Genome-wide FDR = 0.1) than IM and BR considering both years of the study together (Table 2A). Generally, Δ*p_EL_* tends to be larger in magnitude and more variable in Quarry than in IM (Supplemental Figure 2). Across years, 2013 exhibits many more significant Δ*p_EL_* than 2014 (~2-3 fold reduction in 2014). These tests depend on the bulk-specific variance for each sample reported in Table 2B. If each plant in each bulk contributed equally to the DNA library, then ***v*** = 0.005. Several samples are only slightly elevated from this value (e.g. IM, 2014 Early), but the inflation evident in other samples (e.g. BR, 2013 Late) indicate substantial differential representation of sampled genomes in the pool of sequence-suitable DNA.

**Figure 2.**
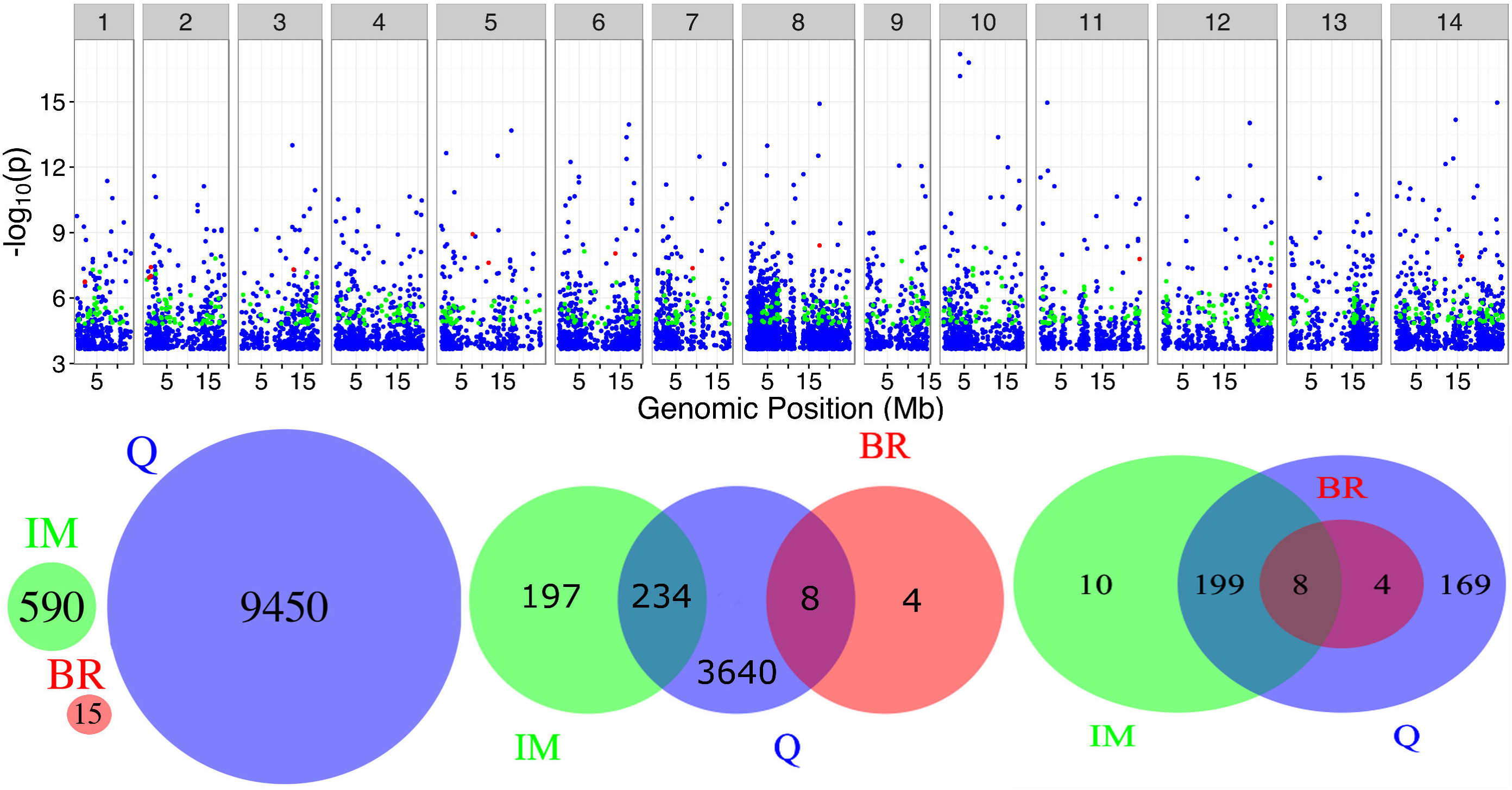
Top: −log_10_(p) for significant Δ*p_EL_* test (FDR=0.1) for each population in 2013. Blue = Quarry, Green = Iron Mountain, and Red = Browder Ridge. Bottom: Overlap in significance across populations in 2013. Left to right: individual sites, 30 kb windows, and 1 Mb windows. A window was considered to be in common between two populations if both populations possessed at least one significant site within the window boundaries.

**Table 2.** (A) Number of significant sites for the individual *Δp_EL_* tests. (B) The estimates for *v*, the bulk-specific variance that aggregates the sampling events prior to sequencing, for each sample.

| (A) | # Significant Tests |  | # Total Tests |  |
| --- | --- | --- | --- | --- |
|  | 2013 | 2014 | 2013 | 2014 |
| IM | 590 | 247 | 3798948 | 4488418 |
| Q | 9450 | 3086 | 4248012 | 5089848 |
| BR | 14 | -- | 3715897 | -- |

**Table 2. (A) Number of significant sites for the individual $\Delta p_{EL}$ tests. (B) The** 5 **estimates for $v$ , the bulk-specific variance that aggregates the sampling events prior** 6 **to sequencing, for each sample.**
| (B) | Browder Ridge | Iron Mountain | Quarry |
| --- | --- | --- | --- |
| 2013 Early | 0.0323 | 0.0141 | 0.0153 |
| 2013 Late | 0.0355 | 0.0200 | 0.0212 |
| 2014 Early | 0.0165 | 0.0066 | 0.0083 |
| 2014 Late | -- | 0.0110 | 0.0130 |

Figure 2 (top) shows the genome-wide distribution of significant sites for each of the three populations from 2013. Here, we observe no overlap of significant SNPs among the different populations (Figure 2, Bottom). However, when we divide the genome into 30 kb or 1 Mb windows, we find that these significant sites are often found in common regions. At both scales, Quarry shares many more significant regions with IM and BR than the latter share with each other. In 2014, there is more overlap despite fewer significant tests (8 SNPs were shared between IM and Quarry). Furthermore, we find few sites to be significant across years within populations: 9 for Quarry and 2 for IM.

The lack of overlap among populations is partially due to significant SNPs in Quarry that are not segregating in the other populations. The heterogeneity/interaction test of Table 1 is limited to SNPs passing filter in multiple samples (in both populations for a given year or in both years for a given population). Results for the three distinct tests (marginal effect, heterogeneity, overall) across the four different contexts are reported in Table 3. As expected, the two contexts displaying the strongest evidence for significant *Δp_EL_* are Quarry and 2013. However, the relative proportion of sites that exhibit a marginal effect versus an interaction effect varies greatly with context. In IM, a nearly equal number of sites are significant for marginal and interaction tests, while in Quarry the vast majority of significant tests are for marginal effects. Similarly, there are relatively few interactions across the two populations in 2014, whereas 2013 is characterized by an almost equal number of SNPs with variable *Δp_EL_*. Importantly, significance for the marginal effect test (M1 v M2) should not be interpreted to mean genuinely fixed effects. Figure 1 indicates that QTL with variable effects can inflate the test statistic for marginal effects (oftentimes more than the heterogeneity test) if *Δp_EL_* has the same direction in each sample.

**Table 3.**
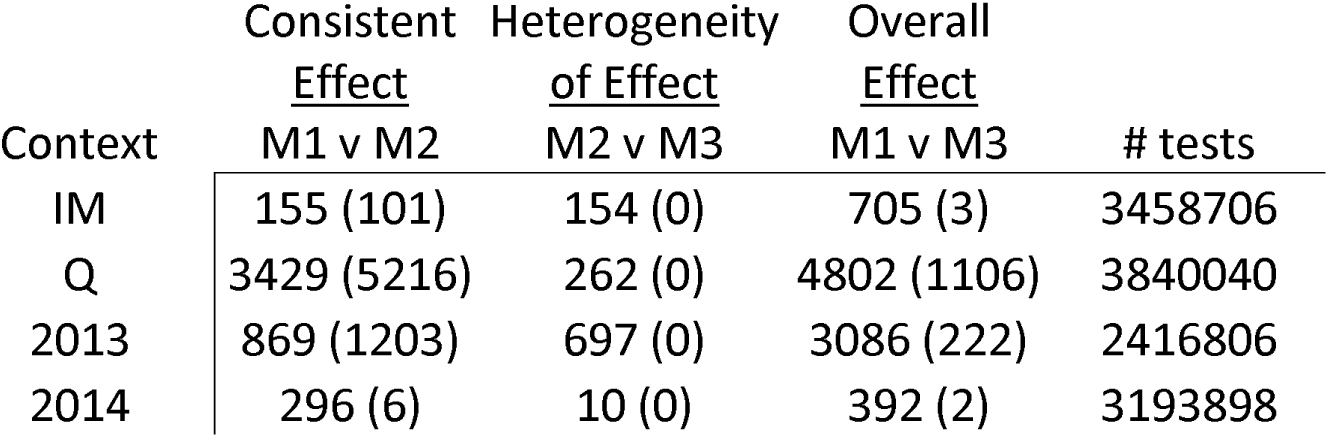
A summary of significance testing for the models of Table 1 is reported for each context. The contrasts are across years within IM and Quarry and across populations in 2013 and 2014, respectively. The number of genome-wide significant tests is reported for both the real data followed (in parentheses) by those obtained from simulation. Simulations were conducted assuming consistent genotypic effects (*a*) and generated with best-matching values of *f_0_* and *c*.

| Context | Consistent<br><u>Effect</u> | Heterogeneity<br><u>of Effect</u> | Overall<br><u>Effect</u> | # tests |
| --- | --- | --- | --- | --- |
|  | M1 v M2 | M2 v M3 | M1 v M3 |  |
| IM | 155 (101) | 154 (0) | 705 (3) | 3458706 |
| Q | 3429 (5216) | 262 (0) | 4802 (1106) | 3840040 |
| 2013 | 869 (1203) | 697 (0) | 3086 (222) | 2416806 |
| 2014 | 296 (6) | 10 (0) | 392 (2) | 3193898 |

### Simulations of loci with consistent effects

To evaluate the results of Table 3 and other features of the data, we calibrated a model of consistent QTL effects for each context (see Supplemental Table 2 for summary of best matching parameter sets; top match was used for simulations). Testing on these simulated data generates a comparable number of significant tests for marginal effects (M1 vs M2): 4749 for real, 6526 for simulated, across contexts (values in parentheses in Table 3). However, the constant effect models are otherwise generally ***inconsistent*** with the real data. The simulations never produced (genome-wide) significant heterogeneity tests (no false positives), but they were abundant in the real data. Additionally, the number of significant outcomes in the overall test (M1 vs M3) was invariably far less than for the marginal test in the simulations, but the opposite is true in the real data (recall that the overall test incorporates signal from both marginal and interaction tests). These discrepancies between simulated and real data indicate genuine variability in Δ*p_EL_* at shared SNPs, particularly across years within IM and across populations within 2013.

Further evidence comes from the covariance of *Δp_EL_* across samples (Figure 3; Supplemental Figures 3 – 6). If genetic effects are constant, this covariance should be substantially positive. Sampling error in estimates for Δ*p_EL_* will reduce the strength of association, but this effect is reiterated in simulations, which are subject to the same degree of sampling variance in Δ*p_EL_*. In all contexts, simulations using best-matching parameters generated an easily detectable positive correlation between Δ*p_EL_* estimates. The real data does not reiterate this pattern. The most striking difference is seen in the 2013 tests (Figure 3 Left) followed by IM (Supplemental Figure 6), both of which have near-zero slopes for the real data, but a strong positive slope for the simulated data. For 2014 and Quarry, the real data exhibits a noticeable positive correlation, which is in agreement with their preponderance of significant marginal-effect tests. In 2014, the slopes for the real and simulated data are near parallel (Supplemental Figure 4), whereas in Quarry the slope for the simulated data is substantially more positive (Supplemental Figure 5). The covariance in *Δp_EL_* across populations (or years) provides a quantitative measure of QTL of (in)consistency (Figure 3 Right). There is evidence of both consistent and variable Δ*p_EL_* sites, but the relative proportion varies across populations and over time.

**Figure 3.**
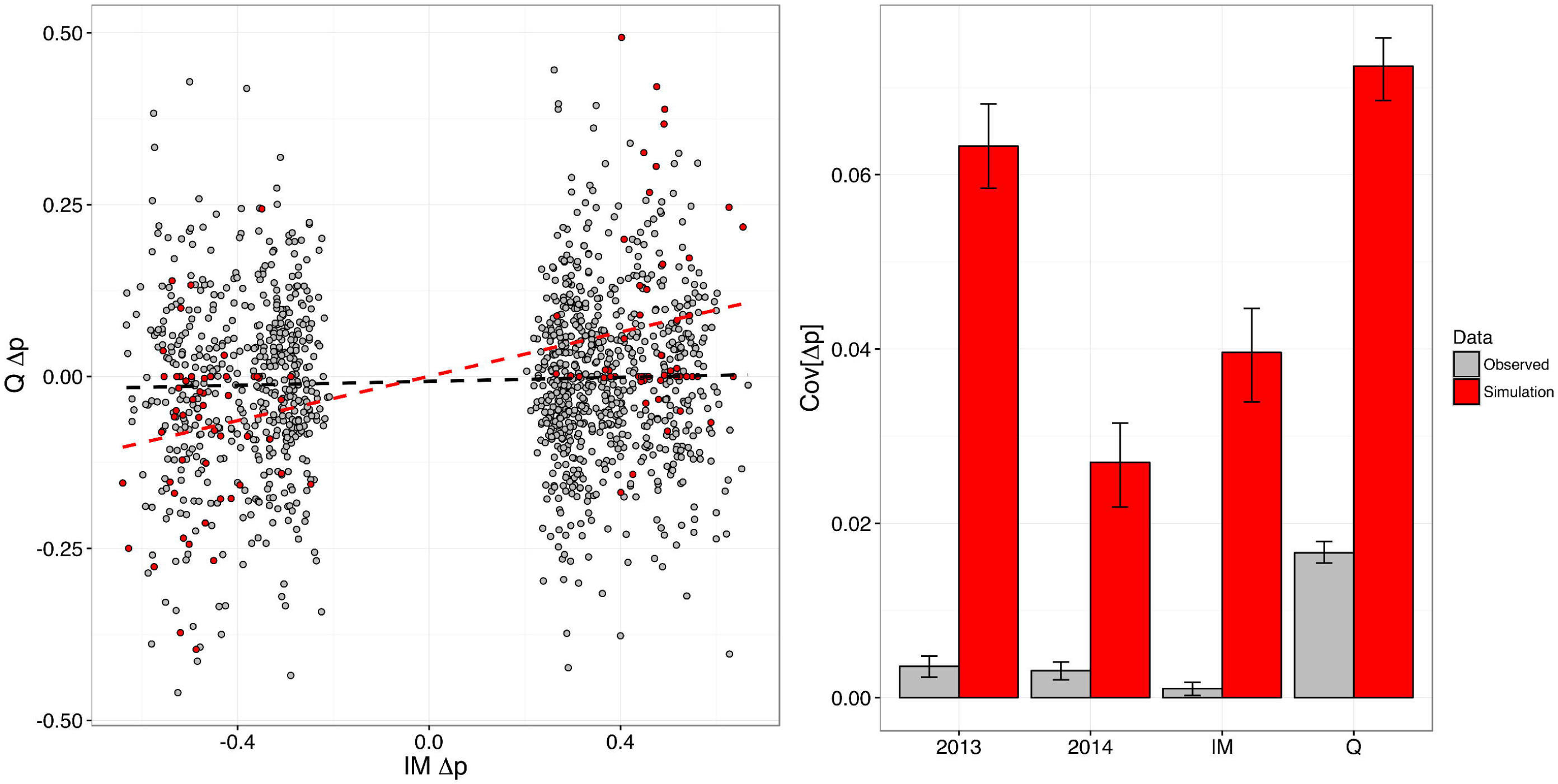
Left: Δ*p_EL_* in Quarry versus IM in 2013 for simulated (red) and real (grey) data. Only sites in which LRT > 15 for the *individual Δp_EL_* in IM are shown. Right: Covariance in Δ*p_EL_* for sites (LRT > 15 for the test of M1 v M3) across time (see bars for IM and Q) and space (2013 and 2014). Confidence intervals are observed 5^th^ and 95^th^ percentiles from bootstrap distribution generated with 100 replicates.

An interesting secondary conclusion from the simulations is that the lack of overlap of significant tests from single Δ*p_EL_* estimates (Figure 2, bottom left) is ***not*** compelling evidence for heterogeneous effects. Even when effects are constant, as implemented in the simulations, shared significance is rare due to an abundance of false negatives. For example, for a pair of populations where consistent effects are relatively frequent and strong (*f_0_* = 0.1 and *a* = 0.3), we found only 353 SNPs to be simultaneously significant out of 26,748 that were deemed significant in either population individually.

### Structural variation

The five structural polymorphisms show strong, but highly variable, effects on flowering time (Figure 4). The first observation is that the *Drive* locus, which was previously known to be polymorphic only in IM and one other population (Case *et al.* 2016), is segregating in Quarry. The *Drive* allele, which enjoys a segregation advantage in female gametes (Fishman and Saunders 2008), is elevated in late flowering samples in IM in both years and in Quarry in 2013. It is enriched in early flowering plants in Quarry in 2014. *inv8*, which had previously been described mainly as a fixed difference between ecotypes, is also segregating in Quarry. We find no evidence for an *inv8* effect in IM, probably because the perennial orientation is rare due to strong local selection (Puzey *et al.* 2016). However, the strong effects in Quarry are consistent with the perennial orientation delaying flowering. Results for *inv6, inv5*, and *inv10* are ambiguous, and it is not clear that the latter two loci are polymorphic in these populations (full results reported in Supplemental Table 4).

**Figure 4.**
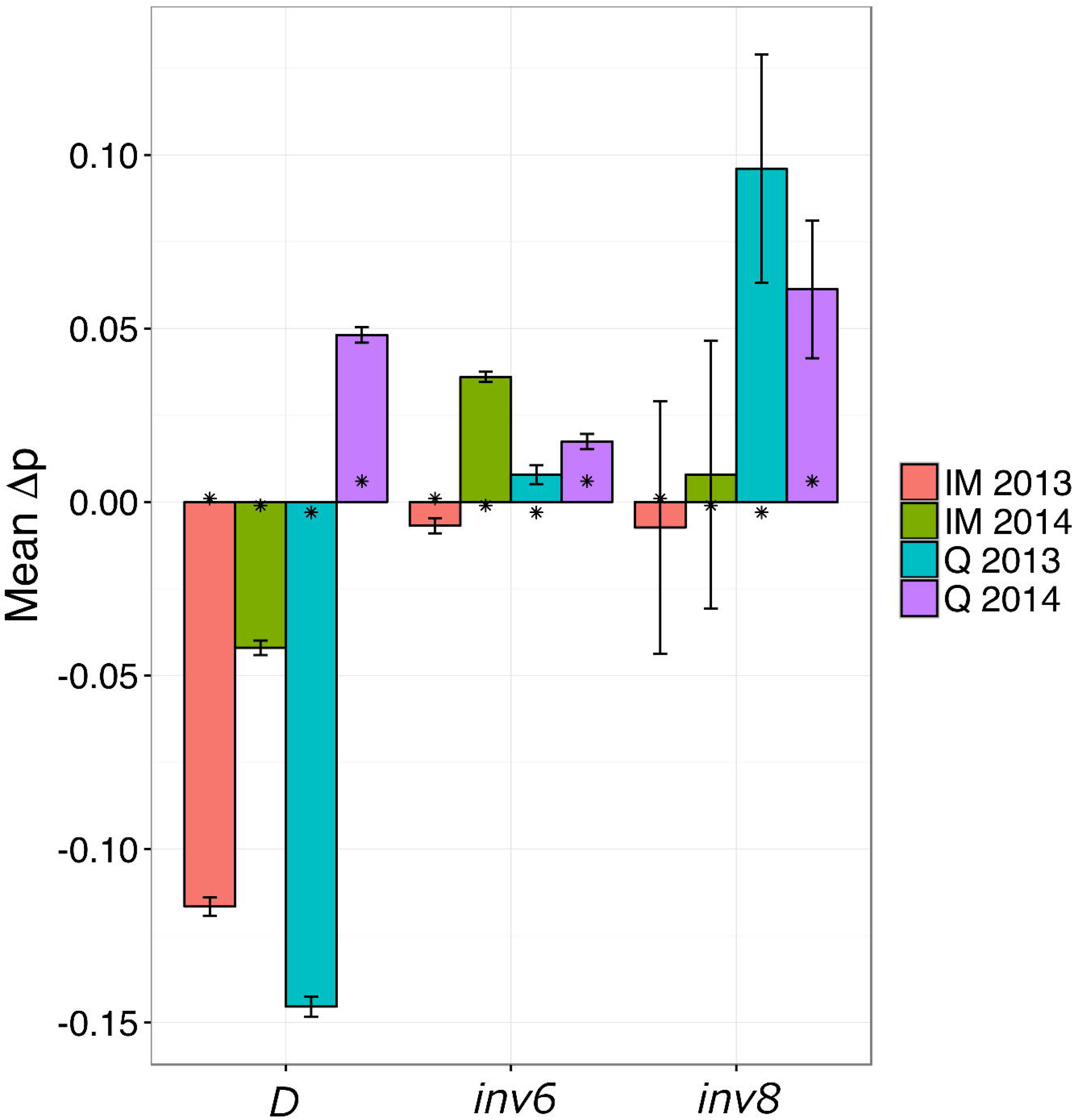
Mean allele frequency divergence between early and late flowering plants for three structural variants. Error bars are +/- SE taken across the SNPs within each feature/population/year.

There is also a clear impact of *inv8* on our SNP-level analyses. In Figure 2, significant Δ*p_EL_* are evenly dispersed with the exception of chromosome 8 (Supplementary Table 3), which has ~5 times more than any other chromosome. This inflation is entirely attributable to *inv8* within Quarry (3281 of the 3,333 significant tests on chromosome 8 are due to Quarry, 2026 of which are within *inv8*). Figure 5 (top) shows a very high density of SNPs significant for the marginal-effect test within *inv8*, and these SNPs are among the highest observed LRTs (see Supplemental Figure 7 for comparison with interaction effect test). Figure 5 (bottom) plots allele frequency over time for the sites with *positive Δp_EL_* (higher reference frequency in early bulk) and in the 99.95 percentile of the LRT for marginal effect in each population. These SNPs produce remarkably consistent oscillations in both IM and Quarry, but are almost entirely non-overlapping (only 4 of the SNPs in Figure 5 and Supplemental Figure 8 are common across populations). This discrepancy is, again, partly due to the presence of *inv8* in Quarry. In Quarry, 306/1921 (15.9%) of the SNPs in the 99.95 marginal-effect LRT percentile are from *inv8*, whereas only 30/1730 (1.7%) are in *inv8* for IM. Also, nearly all of these 306 *inv8* SNPs in Quarry exhibit positive Δ*p_EL_* (281/306 = 91.8%), indicating that the reference (annual) orientation is at higher frequency in early flowering plants. In both populations, there is a tendency towards positive Δ*p_EL_* for these consistent SNPs (918 positive Δ*p_EL_* versus 812 negative Δ*p_EL_* in IM; 1255 positive Δ*p_EL_* versus 666 negative Δ*p_EL_* in Quarry). This tendency is exaggerated in Quarry even after accounting for the effect of *inv8* (974 positive versus 641 negative sites are non-*inv8*). Interestingly, we find that a majority of sites in this 99.95 percentile are at overall high reference frequency in both populations, with many of these sites fixed for the reference allele in either the early or late flowering plants (note the high density of sites in the upper portion of Figure 5 and Supplemental Figure 8).

**Figure 5.**
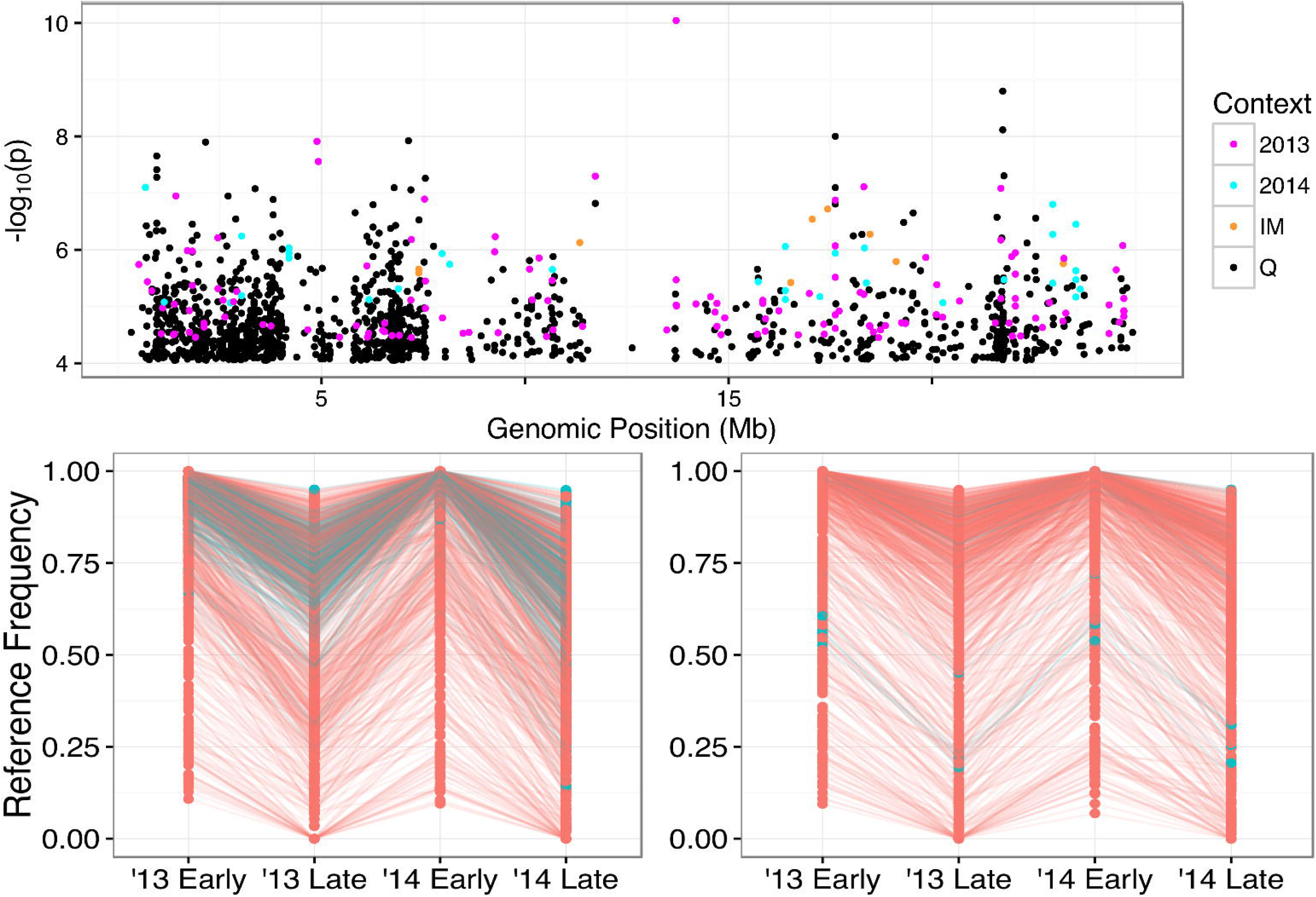
−log_10_(p) for the marginal-effect test plotted along chromosome 8 (Top). On bottom, reference frequency across the four sampling events for sites in the 99.95 percentile of the marginal-effect test that exhibit a *positive Δp_EL_.* Left = Quarry, Right = Iron Mountain. Color indicates whether site is within the chromosome 8 inversion (blue) or not (red).

## Discussion

Our question is the extent to which the loci generating intra-population variation in quantitative traits are consistent within a species. Is the average effect of a QTL similar in neighboring populations, or even in the same population from one generation to the next? We develop a likelihood based testing procedure to distinguish consistent and heterogeneous effects and then apply the procedures to genomic data from ten population samples. Synthesizing multiple aspects of the results across populations and years, the experiment strongly supports heterogeneity of QTL effects. This suggests appreciable lability of allelic effects in nature and underscores the importance of a broad sampling of natural variation in genetic mapping studies. Additionally, the results inform the potential for, or perhaps the expected scale at which, parallel or repeated evolution may occur. In the following sections, we discuss explanations for the observed variation within and among populations and their implications for evolution in nature.

### Why does *Δp_EL_* vary across time and space?

We map loci from the difference in allele frequency between the earliest and latest flowering plants in a population, Δ*p_EL_*. The expected value for *Δp_EL_* depends on allele frequencies, the selection intensity, the phenotypic variance, and the average effect of alleles (equation 6c; Falconer and mackay (1996), pg 200). We controlled selection intensity with our sampling method, but it is clear that each of the other three components varied across space or time in this experiment. Allele frequency differences are clearly important in explaining the differences among populations. Many intermediate frequency SNPs in Quarry that exhibited significant Δ*p_EL_* are fixed (or at least nearly so) within IM and BR. We attribute the elevated genomic and phenotypic variation in Quarry to recent hybridization of annual and perennial genotypes of *M. guttatus* (Monnahan *et al.* 2015). IM and BR are annual populations, and although each has high polymorphism, they produce far fewer significant *Δp_EL_*.

Quarry is about twice as divergent from each annual population as the annuals are from each other: Genome-wide F_ST_ = 0.132 (Quarry versus IM), 0.124 (Quarry versus BR), and 0.065 (BR versus IM; Supplemental Table 3 from (Monnahan *et al.* 2015)). This is important not only because Δ*p_EL_* is proportional to *p(1-p)* at the SNP in question, but also because divergence among populations will also affect the distribution of genomic backgrounds in which that SNP is expressed. Changes in average effect with genomic background (epistasis) has been demonstrated in greenhouse studies of *M. guttatus* for numerous life history traits, including flowering time (Kelly and Mojica 2011; Monnahan and Kelly 2015a; Monnahan and Kelly 2015b).

Differences in the environment must also be important and can alter *Δp_EL_* in several ways. Despite the physical proximity of these populations, they differ dramatically in a number of environmental variables that affect flowering time. Quarry is a south-facing population, IM faces west, and BR faces east; each experiences differing sun exposure. The Quarry site clears of snow earliest in the season (and thus enjoys a longer growth interval) and also has a much shallower grade, particularly in comparison to Iron Mountain. The primary water source for these plants is from snowmelt, and the shallow grade means that water moves slower and perhaps lasts longer for Quarry. Lastly, the edaphic substrate differs greatly between populations; dirt and gravel at Quarry whereas the other two populations grow on a shallow bed of moss atop bedrock. Roots penetrate much deeper at Quarry allowing plants access to additional water and perhaps a different nutrient profile compared to the other populations.

Genotype-by-environment (GxE) interactions are routinely observed in QTL experiments and can be appreciable in magnitude relative to the marginal effect across environments (Scheiner 1993). GxE can change the average effect across populations if there is spatial variation in environmental variables. GxE is the most likely cause of temporal heterogeneity in Δ*p_EL_* (e.g. IM in Table 3), because other factors such as differences in allele frequency (and thus the distribution of genetic backgrounds) should be relatively limited between successive generations within a population. A major temporal fluctuation between the two years of this study was time of snow melt (i.e. beginning of the growing season). Snow cleared in May of 2013, but as early as mid-March in 2014. There was also a late bout of rain in mid-July 2014 extending an already elongated growing season. Furthermore, epistasis and GxE may themselves interact. Significant three-way interactions (GxGxE) have been documented in both field and laboratory studies (Caicedo *et al.* 2004; Zhu *et al.* 2014; Joseph *et al.* 2015; Monnahan and Kelly 2015b).

In addition to GxE, environmental variation can alter Δ*p_EL_* via at least two other routes. First, the predicted Δ*p_EL_* is inversely proportional to the phenotypic standard deviation of the trait. Thus, a shift in environmental conditions that increases the environmental component of variation will reduce Δ*p_EL_*, all else equal. Consistent with this effect, the phenotypic variance in flowering time was elevated in 2014 relative to 2013 (a greater number of days accrued between early and late collections in both IM and Quarry) while the number of significant Δ*p_EL_* tests was reduced (2014 count less than half the 2013 count across IM and Quarry). A second effect of environmental variation on *Δp_EL_* is indirect. Sustained spatial heterogeneity in environmental variables will generate divergent selection and consequent local adaptation. This may be a major cause of allele frequency differences among populations, which subsequently generate differences in Δ*p_EL_*.

The context with the greatest consistency of effects was between years in Quarry (Table 3; Figure 3), which may be attributable in part to the hybrid nature of this population. This population was established no more than 40 generations ago when a rock quarry fell into disuse and was subsequently colonized by nearby *M. guttatus.* Extensive linkage disequilibrium (LD) confirm that the population remains highly admixed (Monnahan *et al.* 2015), which likely reflects both recent formation and continued immigration. Nearly all polymorphic SNPs in IM and BR also segregate in Quarry, but the reverse is not true. Alternative “alleles” may be fairly substantial haplotypes; descendent from annual or perennial ancestors (or immigrants). Such alleles will be “large-effect” if they aggregate the effects of numerous linked polymorphisms.

The average Δ*p_EL_* in this context should be larger relative to estimation error, increasing the number of significant tests for a marginal effect (Table 3), and positive, given the annual nature of the reference genome and typically delayed flowering in perennials (see final paragraph in *Results*). The high LD should also inflate the number of non-causal SNPs exhibiting significant Δ*p_EL_*, hitch-hikers in the terminology of (Maynard Smith and Haigh 1974). While Quarry has greater actual genetic variation in flowering time, LD should further inflate the number of significant tests. LD could also exaggerate our observation in the temporal consistency of Δ*p_EL_* across years (Figure 5).

The results from Quarry underscore a number of general points about the analysis. First, while there are thousands of significant SNPs across populations/years, the number of functionally important variants is likely much smaller. A causal locus for flowering time will “pull” on neighboring SNPs in LD; an effect most pronounced in Quarry but not negligible in IM or BR. Whole genome sequencing of lines from IM indicates substantial LD among SNPs at the gene level (inter-SNP distances of hundreds to a few thousand bp; (Puzey *et al.* 2016)). In principle, assortative mating owing to differences in flowering time might generate substructure within populations. If strong enough, such structure might allow LD among unlinked SNPs. In the present study, we do not find strong internal structure. Divergence measured as Fst is much lower between Early and Late flowering plants within populations (about 1-2%) than it is between populations (12-13% between Quarry and IM or BR).

### Measuring effects for a highly polygenic trait

A recent study of body pigmentation in fruit flies provides a striking contrast to our results. Endler *et al.* (2016) compared populations of *Drosophila melanogaster* from Europe and South Africa using a similar bulked-segregant approach. In contrast to the results here, they found relatively consistent architecture across populations. Genome-wide significant tests were contained within two genic regions, both shared between Europe and South Africa. One important difference is that Endler *et al.* (2016) measured phenotypes from animals reared under common laboratory conditions, thus limiting GxE interactions. A second critical difference is the nature of the traits under study. Coloration phenotypes in both plants and animals are frequently (although not always) influenced by a few major factors (Epperson and Clegg 1988; Joron *et al.* 2006; Steiner *et al.* 2007; Smith and Rausher 2011). In contrast, flowering time is a highly polygenic trait with extensive environmental influence (Coupland 1995; Simpson and Dean 2002).

The best examples of “major loci” in the present study are the structural polymorphisms segregating in IM and Quarry (Figure 4). Our *Δp_EL_* estimates at these loci are, in one sense, likely the most precise in the experiment because each is based on an average across many SNPs. This averaging should minimize estimation error due to finite sequence depth, although not due to the finite sampling of individuals into bulks. Admittedly, we are likely underestimating the magnitude of Δ*p_EL_* for the inversions owing to imperfect association between “diagnostic” SNPs and the actual alternative alleles (inversion orientations). Assuming underestimation to be minor, and noting that the additive variance contributed by a QTL is *2p(1-p)a^2^* (Falconer and Mackay 1996), we use observed *Δp_EL_* values to estimate the variance contribution of QTL. An observed *Δp_EL_* of 0.1 (like *inv8* in Quarry, 2013) is predicted for a locus that explains almost but not quite 1% of the phenotypic variance. If *Δp_EL_* = 0.15 (like the Drive locus in Quarry, 2013), the locus explains 1.5% of the phenotypic variance. While these calculations are coarse, they do emphasize that major flower time loci are decidedly quantitative in their effects.

Many of our estimates for *Δp_EL_* at significant SNPs are large in magnitude (greater than 0.4; Figure 3, left). However, when considering single SNPs, it is essential to recognize that magnitude is inevitably overestimated in the pool of significant tests (Beavis 1994; Ioannidis 2008). For this reason, the simulation study is fundamental to our conclusion of genuine heterogeneity in the effects of flowering time loci. Our simulations reiterate the stochastic processes generating exaggerated values for *Δp_EL_* and, also, the ascertainment process by which overestimated values are used for subsequent analyses. These factors are clearly important. For example, the association of Δ*p_EL_* between two populations with data generated from constant effect loci is positive (red line of Figure 3), but the slope is greatly reduced from 1, which would be the slope of the regression if there were no estimation error. It is the fact that the covariance of *Δp_EL_* between samples within each context is significantly lower than predicted (Figure 3, right), ***after accounting for error and ascertainment***, which indicates heterogeneity.

Our SNP-level hypothesis testing framework was developed to address two basic issues. The first was to provide statistical evidence regarding the marginal effects of QTLs (averaged over populations) as well as the heterogeneity of effects (across populations). The second issue is proper accounting for multiple sources of error inherent to serial sampling in Poolseq studies. Despite best efforts in DNA quantification, pipetting, etc., variable representation of individuals among the sequenced reads is unavoidable. Contingency tables based directly on read counts (e.g. chi-square, Fisher’s exact test) ignore all sampling events prior to the last; essentially treating each read as an independent draw from the ancestral population. Supplemental Table 5 illustrates that contingency table tests can be substantially anti-conservative with respect to our method, at least when the bulk specific sampling variance is non-trivial. Tail p-values can be 1000-fold lower using Fisher’s exact test or chi-square. However, the comparisons also indicate that our likelihood ratio test can occasionally produce lower p-values than the table analyses if the average allele frequency is close to 0 or 1. Such SNPs will usually not be genome-wide significant because there is limited scope for differences in allele frequency between samples if the average is close to 0 or 1. Still, this observation reminds that the arcsin square-root transform does not generate true normality; it just affords a better approximation.

### Flowering time loci

Two qualitatively different kinds of loci are investigated in this experiment. The first are structural polymorphisms, previously mapped in *M. guttatus* although not necessarily known from these populations. The second are SNPs outside of these regions in (presumably) freely recombining parts of the genome. While the strength of evidence for flowering time effects of this latter class may be weaker, they potentially provide much finer resolution. For the structural variants, we cannot distinguish the effects of polymorphisms across the hundreds of genes within each inversion. For other significant SNPs, we located each in relation to putative flowering time genes (based on *M. guttatus* v2 genome annotation; https://www.phytozome.jgi.doe.gov). We considered all SNPs significant for the M1 v M3 test in the genic region of a candidate gene or ±2 kb of the flanking DNA.

In Quarry, 46 significant SNPs were located to flowering time genes. These include genes from the photoperiod pathway and gibberellic acid pathway, as well as multiple interacting genes within each pathway. Gibberellic acid has direct effects on floral development, but also indirectly influences flowering time via its effects on germination and general growth regulation (Mouradov *et al.* 2002). Seven of the 12 candidates in this pathway are gibberellin oxygenases, which generally degrade GA and its precursors (Wuddineh *et al.* 2015). Three of these (*Migut.M00902, Migut.M00908*, and *Migut.M00909)* are on a 50 kb stretch of chromosome 13 and all show highly consistent *Δp_EL_* across years ( Δ*ρ̄*_13_ *=* –0.36 and Δ*ρ̄*_14_ = –0.24). Interestingly, two of these genes (*Migut.M00908* and *Migut.M00909*) were also identified in IM and exhibit a similar pattern across years (Δ*ρ̄*_13_ = –0.44 and Δ*ρ̄*_14_ = –0.26). In addition, both Quarry and IM identified GAI *(Migut.H01666*) as a candidate, a transcription factor that represses GA responses (PENG *et al.* 1997). In aggregate, these results support the GA pathway as a general source of natural variation in flowering time.

Critical photoperiod requirements are typically much longer for perennial *M. guttatus*, with most perennials (and even some annuals) requiring vernalization upon previous exposure to short-day conditions (Friedman and Willis 2013). In Quarry, a SNP ~1.5 kb downstream of VERNALIZATION1 (VRN1; *Migut.H02193*) shows a consistent difference across years (Δ*p*_13_ = 0.54 and Δ*p*_14_ = 0.27). This SNP is within the major photoperiod and vernalization QTL mapped by Friedman and Willis (2013). While VRN1 is a “vernalization” gene, it is transcription-responsive to photoperiod (Dubcovsky *et al.* 2006) and has distinct effects on flowering time apart from vernalization (Levy *et al.* 2002). ELF6 (Early Flowering; *Migut.F01729*) (Clouse 2008), a repressor of the photoperiod pathway, also shows consistently higher reference base frequency in the early flowering samples (Δ*p*_13_ = 0.29 and Δ*p*_14_ = 0.20; significant for both the M1 v M2 and M1 v M3 tests). Significant SNPs were also found adjacent to ELF 3 and ELF4 (*Migut.E01551* and *Migut.J00944*, respectively), and again, the reference base frequency was higher in early flowering plants. However, *Δp_EL_* was less consistent across years (ELF3: Δ*p*_13_ = –0.07 and Δ*p*_14_ = 0.38; ELF4: Δ*p*_13_ = 0.55 and Δ*p*_14_ = 0.04). The direction of these differences (*Δp_EL_* usually positive) may reflect the fact that the reference genome is based on an annual genotype. Thus, the reference base is more likely to be the “annual” allele in an annual/perennial population. Lastly, GIGANTEA (*Migut.C00380*), a major photoperiod response regulator that interacts with multiple ELF transcription factors (Mishra AND Panigrahi 2015), exhibited a significant interaction across years in Quarry, with Δ*p*_13_ =–0.25 and Δ*p*_14_ = 0.14.

Both IM and Q also have significant SNPs in a tandem pair of GDSL-motif lipase genes (*Migut.M01081* and *Migut.M01082*) as well as an RNA ligase gene (*Migut.N02091*). The former belong to a class of lipases with broad, ecologically-relevant functions including microbial defense (Oh *et al.* 2005; Kwon *et al.* 2009), morphogenesis and development (Riemann *et al.* 2007; Lee *et al.* 2009), and abiotic stress responses (Hong *et al.* 2007). While these genes may play a direct role in promoting/hindering development, it may just as well function in defense of a pest associated with early or late season conditions. RNA-ligase is involved in the maturation of tRNAs, and it was recently discovered that mutants of an RNA-ligase gene in *Arabidopsis thaliana* exhibited defects, specifically, in auxin-related growth processes (Leitner *et al.* 2015).

The structural polymorphisms provide clear evidence of flowering time effects (Figure 4), although without gene-level resolution. However, the estimates for phenotypic and fitness effects for entire karyotypes is valuable when considering the evolutionary dynamics of these polymorphisms. The results for *inv8* are fully consistent with expectations based on previous studies of this locus. Alternative orientations distinguish annual and perennial ecotypes of *M. guttatus* and QTL mapping reveals large effects of *inv8* on flowering time, anthocyanin production, and growth-related traits (Lowry and Willis 2010). As in the mapping study, our experiment shows that the perennial orientation delays flowering, and its presence confirms the annual/perennial origin of this population. This study provides the most direct evidence of *inv8* segregating within a natural population, contributing to phenotypic variation; although polymorphism in other populations is suggested (Twyford and Friedman 2015).

The strong effect of the meiotic drive locus on flowering time is more surprising. However, field experiments have demonstrated *Drive* effects on both male and female fitness components (Fishman and Kelly 2015), which may depend on flowering time. Direct effects of this locus on developmental timing have been documented in a greenhouse experiment (Scoville *et al.* 2009). It is possible that some delay in flowering is due to the reduction in pollen viability caused by the *Drive* karyotype. Bee pollinators discriminate against flowers with lower viable pollen (Carr *et al.* 2014) and lack of visitation prolongs flower lifespan (Arathi *et al.* 2002). Finally, the derived orientation of *inv6* was associated with earlier flowering in IM in 2014, but not the previous year (Figure 4). Several greenhouse studies have shown *inv6* effects on days to flower (Lee 2009; Scoville *et al.* 2009) although the direction of effect varies with genetic background and perhaps the sequence of the ancestral orientation (which is highly variable and different among experiments).

### Conclusion

We have developed and implemented a method to map genomic regions affecting ecologically-relevant traits directly within natural populations, while accounting for estimation error in observed allele frequencies. Replicated comparative mapping can inform fundamental biological questions such as how the evolutionary trajectories of local populations will transform an entire species. Uniform selection across a species range generated by climate change might set the stage for parallel evolution, but at what scale will parallelism occur? As sequencing costs continue to decrease, the number of populations and range of distribution that can be surveyed will increase. Though our study focuses on a narrow geographic range, it provides a baseline understanding for how genomic variation in flowering time varies across neighboring populations and from generation to generation.

While most flowering time loci varied between populations and over time, a subset exhibited fairly consistent effects. These consistent loci, which include large structural variants, such as inversions, and genes in known flowering time pathways, are most likely to evolve in parallel if populations were to experience uniform selection on flowering time. The actual degree of parallelism will depend on genetic factors (e.g. the distribution of additive and dominance effects), demographic factors (e.g. population sizes, growth rates, and migration), and selective factors (e.g. strength and consistency of selection on flowering time). However, the existence of consistent loci supports the growing body of evidence that parallel evolution can occur from the recruitment of standing genetic variation (Pigeon *et al.* 1997; Colosimo *et al.* 2005; Jones *et al.* 2012). Statistical considerations aside, this consistency also supports the utility of mapping studies to identify a subset of loci that are general contributors to natural variation within and between populations.

In contrast, the observed *variation* in genomic architecture testifies to the influence of divergent environmental conditions and genomic backgrounds on the average effects exhibited by segregating variants. For highly polygenic traits such as flowering time, we would almost certainly expect to find some loci to have evolved in parallel, but this would likely account for a relatively minor portion of the total selection response. Furthermore, a lack of parallelism at the genetic level would not imply that populations did not have access to the same standing variation and thus evolutionary trajectories. Rather, it could simply reflect the idiosyncratic interplay between the factors outlined above. Additional studies will help determine whether variation in genomic architecture is a trait- or species-specific phenomenon as well as highlight those genes that are consistently important drivers for natural variation.

## Supplemental Material

**Supplemental Appendix 1.** Description of estimation of v terms as well as their sampling variance.

**Supplemental Table 1.** Collection dates for each sampling event

**Supplemental Table 2.** Parameter values for simulated data for each of the four test types: IM, Q, 2013, and 2014, as well as their fit to the observed distributions likelihood ratio tests comparing Model 1 versus Model 2. Values for Pr[X>10] and Pr[X>15] for the real data are: 0.0043186 and 0.00053546 for IM, 0.0062383 and 0.0010148 for Q, 0.0052888 and 0.00074395 for 2013, and 0.0036069 and 0.000434581 for 2014.

**Supplemental Table 3.** Number of significant sites on each chromosome, as well as the rate of significant sites per Mb. Tally includes all sites regardless of which population/year they were deemed significant in.

**Supplemental Table 4.** Mean allele frequency difference between early and late flowering plants for structural variants.

**Supplemental Table 5.** Comparison of Fisher’s exact test, contingency table analysis, and our likelihood ratio test for read counts from chromosome 1 in Browder Ridge. Results contained in separate spreadsheet.

**Supplemental Figure 1.** Venn diagrams depicting sites passing filter in one or more of the populations Top: 2013, Bottom: 2014.

**Supplemental Figure 2.** Genome-wide averages of *Δp_EL_* and *Δp_EL_*^2^ (proportional to variance), calculated in 1 Mb windows.

**Supplemental Figure 3.** Left: *Δp_EL_* in Quarry versus IM in 2013 for simulated (red) and real (grey) data. Only sites in which LRT > 15 for the *individual Δp_EL_* in IM are shown. Right: Same as left except only sites in which LRT > 15 for the *individual* Δ*p_EL_* in **Q** are shown. Values used for simulation: *f_0_=*0.08 *a*=0.205

**Supplemental Figure 4.** Left: *Δp_EL_* in Quarry versus IM in 2014 for simulated (red) and real (grey) data. Only sites in which LRT > 15 for the *individual Δp_EL_* in IM are shown. Right: Same as left except only sites in which LRT > 15 for the *individual* Δ*p_ΕL_* in **Q** are shown. Values used for simulation: Top)*f_0_*=0.145 *a*=0.115. Bottom) *f_0_=0.*06 *a*=0.16.

**Supplemental Figure 5.** Left: Δ*p_ΕL_* in 2013 versus 2014 in Quarry for simulated (red) and real (grey) data. Only sites in which LRT > 15 for the *individual Δp_EL_* in 2013 are shown. Right: Same as left except only sites in which LRT > 15 for the *individual* Δ*p_ΕL_* in **2014** are shown. Values used for simulation: Top)*f_0_*=0.065 *a*=0.215 Bottom) *f_0_=0.*10 *a*=0.19

**Supplemental Figure 6.** Left: Δ*pEL* 2013 versus 2014 in IM for simulated (red) and real (grey) data. Only sites in which LRT > 15 for the *individual Δp_EL_* in 2013 are shown. Right: Same as left except only sites in which LRT > 15 for the *individual Δp_EL_* in **2014** are shown. Values used for simulation: Top)*f_0_*=0.20 *a*=0.125. Bottom) *f_0_=0.*20 *a*=0.13

**Supplemental Figure 7.** Significant sites (FDR = 0.1) for test of marginal-effect (top) and interaction effect (bottom) for each of the four contexts on chromosome 8.

**Supplemental Figure 8.** Reference frequency across the four sampling events for sites in the 99.95 percentile of the marginal-effect test that exhibit a positive *Δp_EL_* (left; same as Figure 5) and those that exhibit a negative Δ*p_EL_* (right). Top = Quarry, Bottom = Iron Mountain. Color indicates whether site is within the chromosome 8 inversion (blue) or not (red).

## Data Deposition

BioProject PRJNA336318. BioSamples: SAMN05508935, SAMN0550981, SAMN0550982, SAMN0550983, SAMN0550984, SAMN0550985, SAMN0550986, SAMN0550987, SAMN0550988, SAMN0550989, and SAMN0550990.

## Acknowledgements

We would like to thanks the following individuals for the helpful comments on the manuscript: SJ MacDonald, ME Orive, and L Hileman. We acknowledge funding from the KU Botany Endowment, the KU Graduate Research Fund, and National Institute of Health (R01 GM073990-02).

